# Scalable Phylogenetic Profiling using MinHash Uncovers Likely Eukaryotic Sexual Reproduction Genes

**DOI:** 10.1101/852491

**Authors:** David Moi, Laurent Kilchoer, Pablo S. Aguilar, Christophe Dessimoz

**Affiliations:** Department of Computational Biology, University of Lausanne, Switzerland; Center for Integrative Genomics, University of Lausanne, Switzerland; SIB Swiss Institute of Bioinformatics, Lausanne, Switzerland; Instituto de Investigaciones Biotecnologicas (IIBIO), Universidad Nacional de San Martín Buenos Aires, Argentina; Instituto de Fisiología, Biología Molecular y Neurociencias (IFIBYNE-CONICET); Department of Genetics, Evolution, and Environment, University College London, UK; Department of Computer Science, University College London, UK

## Abstract

Phylogenetic profiling is a computational method to predict genes involved in the same biological process by identifying protein families which tend to be jointly lost or retained across the tree of life. Phylogenetic profiling has customarily been more widely used with prokaryotes than eukaryotes, because the method is thought to require many diverse genomes. There are now many eukaryotic genomes available, but these are considerably larger, and typical phylogenetic profiling methods require quadratic time or worse in the number of genes. We introduce a fast, scalable phylogenetic profiling approach entitled HogProf, which leverages hierarchical orthologous groups for the construction of large profiles and locality-sensitive hashing for efficient retrieval of similar profiles. We show that the approach outperforms Enhanced Phylogenetic Tree, a phylogeny-based method, and use the tool to reconstruct networks and query for interactors of the kinetochore complex as well as conserved proteins involved in sexual reproduction: Hap2, Spo11 and Gex1. HogProf enables large-scale phylogenetic profiling across the three domains of life, and will be useful to predict biological pathways among the hundreds of thousands of eukaryotic species that will become available in the coming few years. HogProf is available at https://github.com/DessimozLab/HogProf.

## Introduction

The NCBI Sequence Read Archive (SRA) contains 1.6×10^16^ nucleotide bases of data and the quantity of sequenced organisms keeps growing exponentially. To make sense of all of this new genomic information, annotation pipelines need to overcome speed and accuracy barriers. Even in a well-studied model organism such as *Arabidopsis thaliana*, nearly a quarter of all genes are not annotated with an informative gene ontology term [1]. One way to infer the function of a gene product is to analyse the biological network it is involved in and form a hypothesis based on its physical or regulatory interactors. Unfortunately, biological network inference is mostly limited to model organisms as well and genome scale data is only available through the use of noisy high-throughput experiments.

To ascribe biological functions to these new sequences, most of which originate from non-model organisms, computational methods are essential [reviewed in 2]. Among the computational function prediction techniques that leverage the existing body of experimental data, one important but still underutilised approach in eukaryotes is *phylogenetic profiling [3]*: positively correlated patterns of gene gains and losses across the tree of life are suggestive of genes involved in the same biological pathways.

Phylogenetic profiling has been more commonly performed on prokaryotic genomes than on eukaryotic ones. Perhaps due to the relative paucity of eukaryotic genomes in the 2000s, earlier benchmarking studies observed poorer performance with eukaryotes than with Prokaryotes [4–6]. The situation today is considerably different; the GOLD database [7] tracks over 6000 eukaryotic genomes. Multiple successful applications of phylogenetic profiling in eukaryotes have been published in recent years, e.g. to infer small RNA pathway genes [8], the kinetochore network [9], ciliary genes [10], or homologous recombination repair genes [11].

Still, large-scale phylogenetic profiling with eukaryotes remains computationally challenging, because eukaryotic genomes are larger and more complex than their prokaryotic counterparts, and because state-of-the-art phylogenetic profiling methods typically scale at least quadratically with the number of gene families and linearly with the number of genomes. As a result, most mainstream phylogenomic databases, such as Ensembl [12], EggNOG [13], OrthoDB [14], or OMA [15] do not provide phylogenetic profiles.

The inference of phylogenetic profiles using large datasets is challenging. Some pipelines resort to all-vs-all sequence similarity searches to derive orthologous groups and only count binary presence or absence of a member of each group in a limited number of genomes [16,17] or forego this step altogether and ignore the evolutionary history of each group of homologues, relying instead on co-occurrence in extant genomes [18]. Other tree-based methods infer the underlying evolutionary history from the presence of extant homologues [19]. In our pipeline, we leveraged the already existing OMA orthology inference algorithm, which has been benchmarked and integrates with other proteomic and genomic resources [15]. The OMA database describes the orthology relationships among all protein coding genes of over 2000 cellular organisms. One core object of this database is the Hierarchical Orthologous Group (HOG) [20]. Each HOG contains all of the descendants of a single ancestor gene. When a gene is duplicated during its evolution, the paralogous genes and the descendants of the orthologue are contained in separate subhogs which describe their lineage back to their single ancestor gene (hence the hierarchical descriptor). A brief introductory video tutorial on HOGs is available at https://youtu.be/5p5x5gxzhZA.

Here, we introduce a scalable approach which combines the efficient generation of phylogeny-aware profiles from hierarchical orthologous groups with ultrafast retrieval of similar profiles using locality sensitive hashing. Furthermore, the approach leverages the properties of minhash signatures to allow for the selection of clade subsets and for clade weightings in the construction of profiles. The improvements in performance of our method make it possible to build profiles for the over 2000 genomes contained in OMA. We show that the method is as accurate as a state-of-the-art phylogeny-based method, and illustrate its usefulness by retrieving biologically relevant results for several genes of interest. Because the method is unaffected by the number of genomes included and scales logarithmically with the number of hierarchical orthologous groups added, it will efficiently perform with the exponentially growing number of eukaryotic genomes.

All of the code used to produce the results shown in this manuscript can be downloaded at https://github.com/DessimozLab/HogProf.

## Results

In the following sections we first compare our profiling distance metric against other profile distances in order to characterize the Jaccard hash estimation’s precision and recall characteristics. Following this quantification, we show our pipeline’s capacity in reconstituting a well known interaction network as well as augmenting it with more putative interactors using its search functionality. Finally, to illustrate a typical use case of our tool, we explore a poorly characterized network.

### A scalable phylogenetic profiling method using locality-sensitive hashing and hierarchical orthologous groups

Most phylogenetic profiling methods consist of two steps: creating a profile for each homologous or orthologous group, and comparing profiles. When they were first implemented, profiles were constructed as binary vectors of presence and absence across species [3]. Since then, variants have been proposed, which take continuous values [9]—such as alignment scores with the gene of a reference species [11]—or which count the number of paralogs present in each species. Yet other variants convey the number of events on branches of the species tree [6]. However, all approaches are limited with respect to the computational power required to cluster profiles using their respective distance metrics. Due to this computational cost, profiling efforts are typically focused on reconstructing pathways with known interactors using existing annotations and evidence rather than being used as an exploratory tool to search for new interactors and reconstituting completely unknown networks.

In our approach to the problem of profiling, we captured the evolutionary history of each HOG in enhanced phylogenies and encode them in probabilistic data structures (Fig. 1). These are used to compile searchable databases to allow for the retrieval of coevolving HOGs with similar evolutionary histories and compare the similarity of two HOGs. The two major components of the pipeline that are responsible for constructing the enhanced phylogenies and calculating probabilistic data structures to represent them are pyHam and Datasketch, respectively. Further details on the implementation are provided in the Methods section. The combination of these two tools now allows for the main innovation of our pipeline: the efficient exploration and clustering of profiles to study known and novel biological networks.

**Fig. 1.**
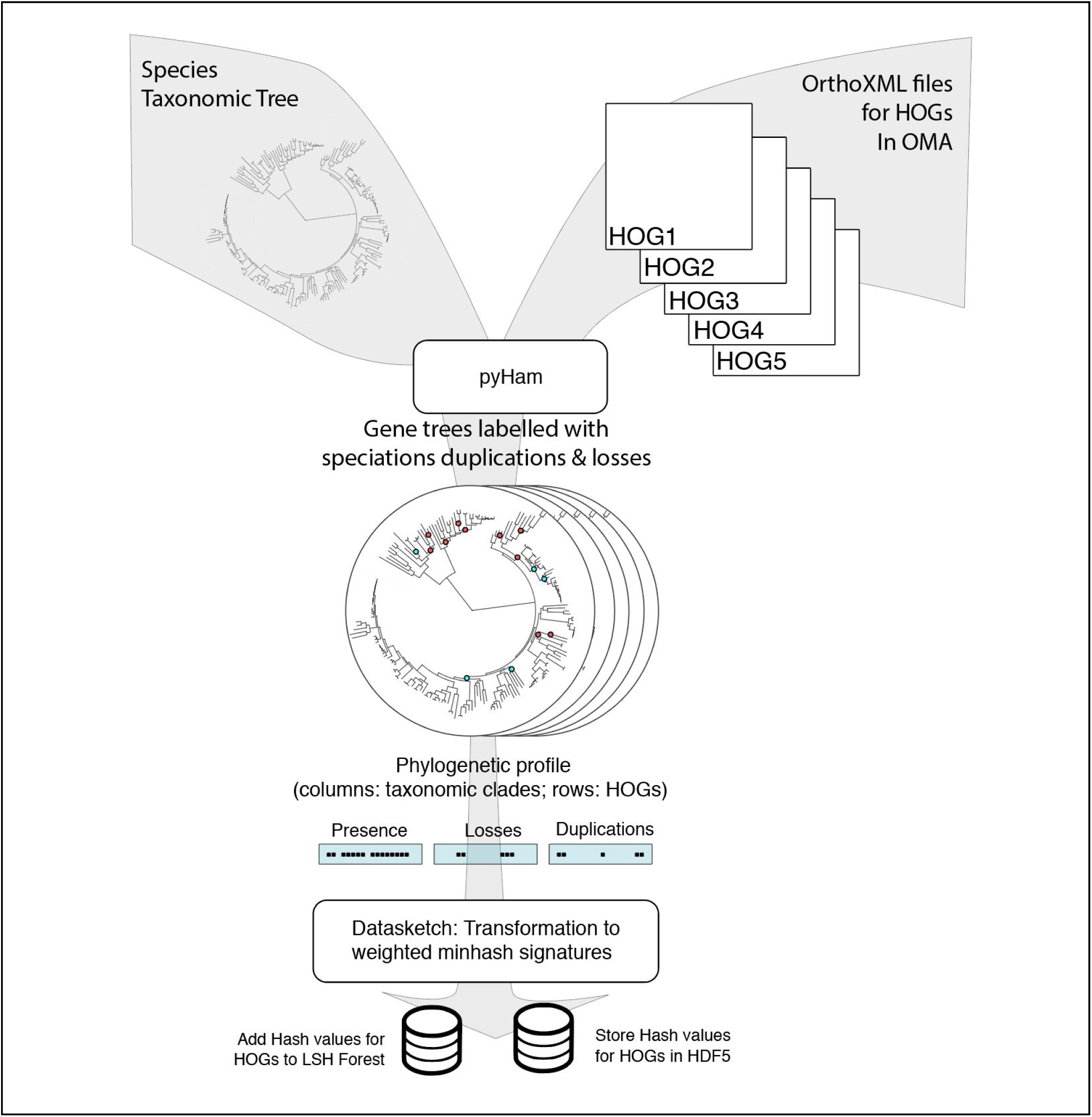
Diagram summarizing the different steps of the pipeline to generate the LSH Forest and hash signatures for each HOG. The labelled phylogenetic trees generated by pyHam are converted into phylogenetic profiles and used to generate a weighted minhash signature with Datasketch. The hash signatures are inserted into the LSH Forest and stored in an HDF5 file.

### Accuracy of predicted phylogenetic profiles in an empirical benchmark

We compared the performance of our profiling metric to existing profile distances using benchmarking data available in Ta *et al.* [16]. In that benchmark, the true positive protein-protein interactions (PPIs) were constructed using data available from CORUM [21] and the MIPS [22] databases for the human and yeast interaction datasets. True negatives were constructed by mixing proteins known to be involved in different complexes. The dataset is balanced with 50% positive and 50% negative samples. Using their Uniprot identifiers, these interaction pairs were mapped to their respective HOGs and their profiles were compared using the hash based Jaccard score estimate. The comparison below shows HogProf alongside other profiling distance metrics that are considerably more computationally intensive, including the Enhanced Phylogenetic Tree metric shown in Ta *et al.* [16]. Yet, our approach outperformed these previous methods, yielding the highest Area Under the Curve for both yeast and human datasets (Figure 2, Table 1).

**Table 1.** AUC values for Profiling distance metrics.

| Metric | AUC Yeast | AUC Human |
| --- | --- | --- |
| Jaccard Hash | 0.6634 | 0.6155 |
| EPT | 0.6104 | 0.5875 |
| BIN PS | 0.5840 | 0.5463 |
| OCC ED | 0.5829 | 0.5268 |
| OCC PS | 0.6028 | 0.5714 |

**Fig. 2.**
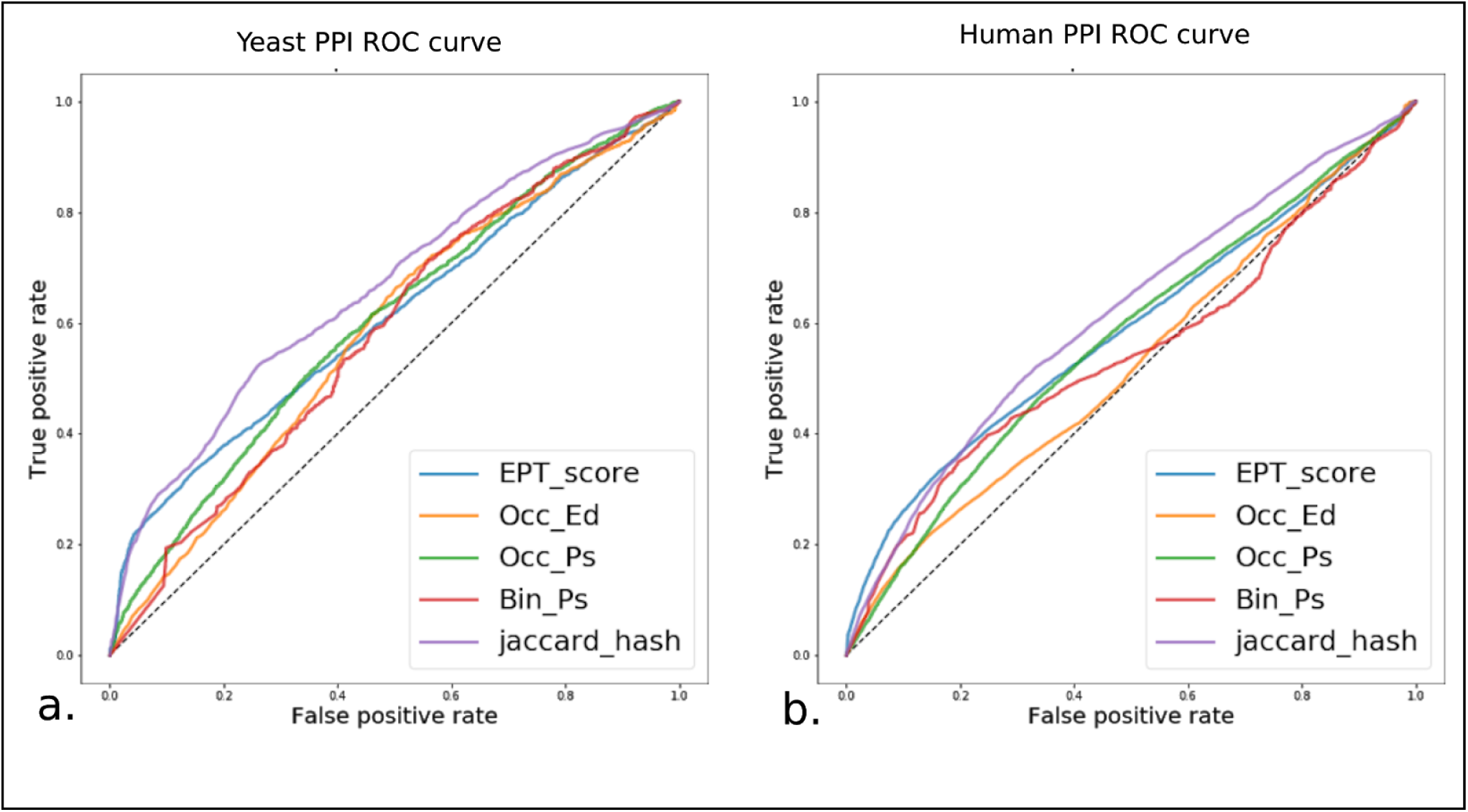
ROC curves for all profiling methods. **a.** Yeast protein-protein interactions. Jaccard Hash and Jaccard Hash Opt perform better than all metrics overall but when high precision is required, EPT score is still slightly more accurate. **b.** Human protein-protein interactions. Jaccard Hash and Jaccard Hash Opt perform better than all metrics overall but again, when high precision is required, EPT score is still slightly more accurate. In both subfigures, Jaccard hash refers to the profiles containing all clades with all weights for each event and taxonomic level set to 1. “EPT_score” refers to the Enhanced Phylogenetic Tree metric developed in [16]. “Bin_Ps” refers to a distance using binary vectors and Pearson correlation described in [23]. “Occ_Ed” and “Occ_Ps” refer to the occurence profiles with Euclidean distance and Pearson correlation as described in [24].

### Recovery of a canonical network: the kinetochore network

To further validate our profiling approach on a known biological network, we used our pipeline to replicate previous work shown in van Hooff et al. [9]. Their analysis focuses on the evolutionary dynamics of the kinetochore complex, a microtubule organizing structure that was present in the last eukaryotic common ancestor (LECA) and has undergone many modifications throughout evolution in each eukaryotic clade where it is found. Its modular organization has allowed for clade-specific additions or deletions of modules to the core complex which remains relatively stable. This modular organisation and clade-specific emergence of certain parts of the complex make it an ideal target for phylogenetic profiling analysis.

We show that our minhash signature comparisons are also capable of recovering the kinetochore complex organisation. After considering just the HOGs for the families used in van Hooff et al. [9], we augmented their set of profiles using the LSH Forest to retrieve interactors that may also be involved in the kinetochore and anaphase promoting complex (APC) network which have not been detected by these authors. By using a well-studied network in eukaryotes to test the LSH Forest search, we can rely on previous work and annotations to quantify the quality of the returned results. Using the Gene Ontology (GO) terms [25] of all proteins returned in our searches for novel interactors, we quantify how enriched they are for the specific functions we would expect to be related to our network of interest.

In their work, van Hooff et al. [9] used pairwise Pearson correlation coefficients between the presence and absence vectors of the various protein components of the kinetochore in the 90 proteomes of a manually selected set of organisms as a distance metric between a manually curated set of profiles corresponding to each component of the complex. Using this pairwise comparison of all vectors, they clustered the profiles and were able to recover known sub-componenents of the complex using just evolutionary information. Using our hash-based Jaccard distance metric in an all-vs-all comparison between the HOGs corresponding to each of these protein families, we were also able to recover the main modules of the kinetochore complex with a similar organisation to the one defined by van Hooff et al. The color clustering in figure 3 corresponds to their original manual definition of these different subcomplex modules based. Despite the vastly different methods used in the construction and comparison of the profiles used to recover the network in both pipelines, we observe that the distance matrices generated by each profiling approach are correlated and are recovering similar evolutionary signals, with Spearman correlation of 0.26 (p < 1e-100) and Pearson correlation of 0.35 (p < 1e-100).

**Fig. 3.**
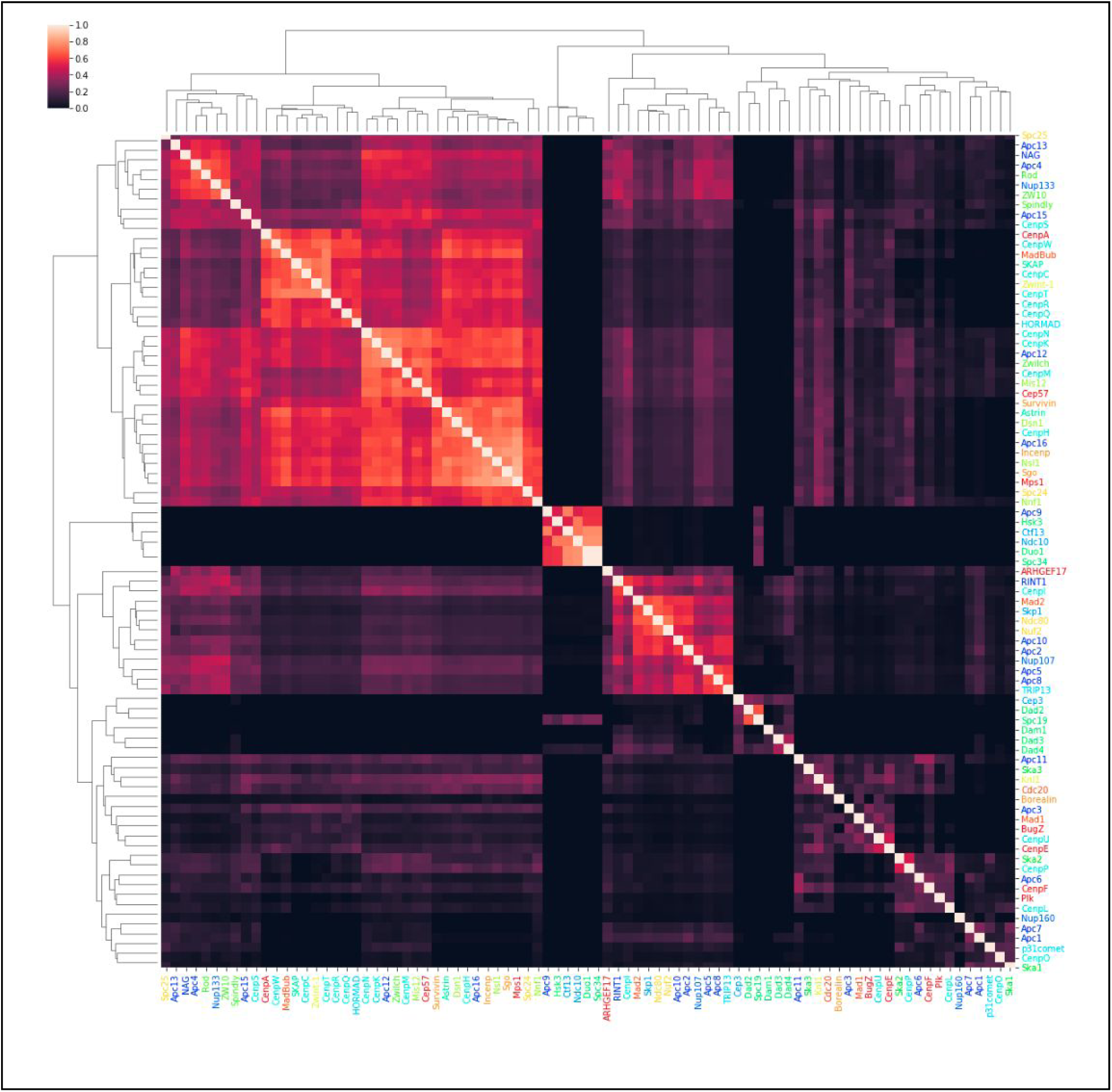
Recovery of kinetochore and APC complexes. After mapping each of the protein families presented in Van Hooff et al. [9] to their corresponding HOG, a distance matrix was constructed by comparing the Jaccard hash distance between profiles using HogProf. Name colors in the rows and columns of the matrix correspond to the kinetochore and APC subcomplex components as defined manually using literature sources in van Hooff et al. All code used to construct the figure is available on the HogProf repository.

The All-vs-All comparison of the profiles reveals several well defined clusters in both works including the Dam-Dad-Spc19 and CenP subcomplexes. However, our profiles were constructed alongside all other HOGs in OMA and were not curated before being compared. With only the information of which proteins were in the complex, we mapped them to their corresponding OMA HOGs and, with this example, demonstrated the ability to reconstruct any network of interest or construct putative networks using the search functionality of our pipeline with minimal computing time. However, it should be noted that the quality of the OMA HOGs used to construct the enhanced phylogenies and hash signatures directly influences our ability to recover complex organisation.

To illustrate the utility of the search functionality of our tool we used the profiles known to be associated with the kinetochore complex to search for other interactors. All HOGs corresponding to the protein families used to analyse the kinetochore evolutionary dynamics in van Hooff et al. [9] were used as queries against an LSH Forest containing all HOGs in OMA. By performing an all-vs-all comparison of the minhash signatures of the queries and returned results, a Jaccard distance matrix was generated showing potential functional modules associated with each known component of the kinetochore and APC complexes.

To verify that the results returned by our search were not spurious, we performed GO enrichment analysis of the returned HOGs that were not part of the original set of queries but appeared to be coevolving closely with known kinetochore components. Given the incomplete nature of Gene Ontology annotations [“open world assumption”, 26], many of these proteins may actually be involved in the kinetochore interaction network but this biological function could be still undiscovered. However, even with this limitation, salient annotations relevant to the kinetochore network were returned in the search results (Table 2 and Supplementary Data 1). The identifiers of all protein sequences contained in the HOGs returned by the search results were compiled and the GO enrichment of each cluster shown in Figure 4 was calculated using the OMA annotation corpus as a background. The enrichment results were manually parsed and salient annotations related to HOGs were selected to be reviewed further in the associated literature to check for the association of the search result with the query HOG (Table 2).

**Table 2.**
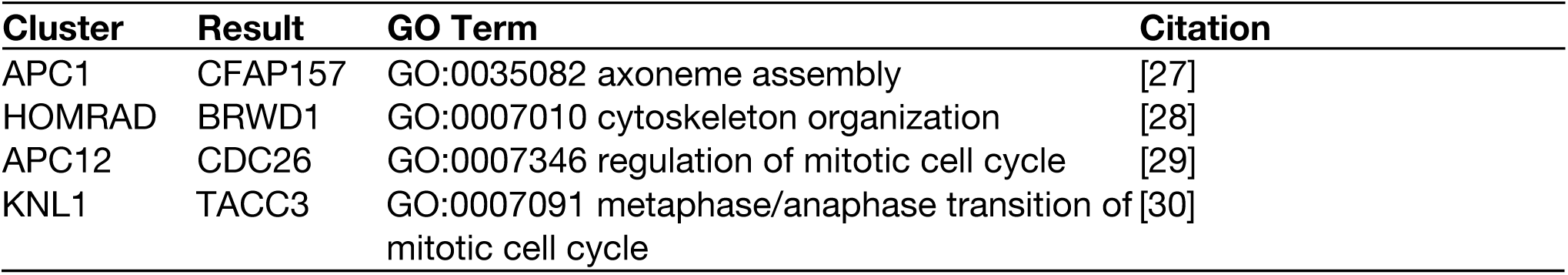
Manually curated biologically relevant search results for interactors coevolving with van Hooff *et al.*’s kinetochore and APC selected protein families[9]. Notable protein families (Result) returned within clusters containing query HOGs (Cluster) are listed with their pertinent annotation and literature. GO enrichment results of clusters that contained one or more queries from the original kinetochore network were analyzed manually. We searched for literature supporting the relevant GO annotations, thereby confirming that the results returned by the LSH were associated with kinetochore and APC processes. This is a non-exhaustive summary of some selected results. The full enrichment results are available as Supplementary Data 1.

**Fig. 4.**
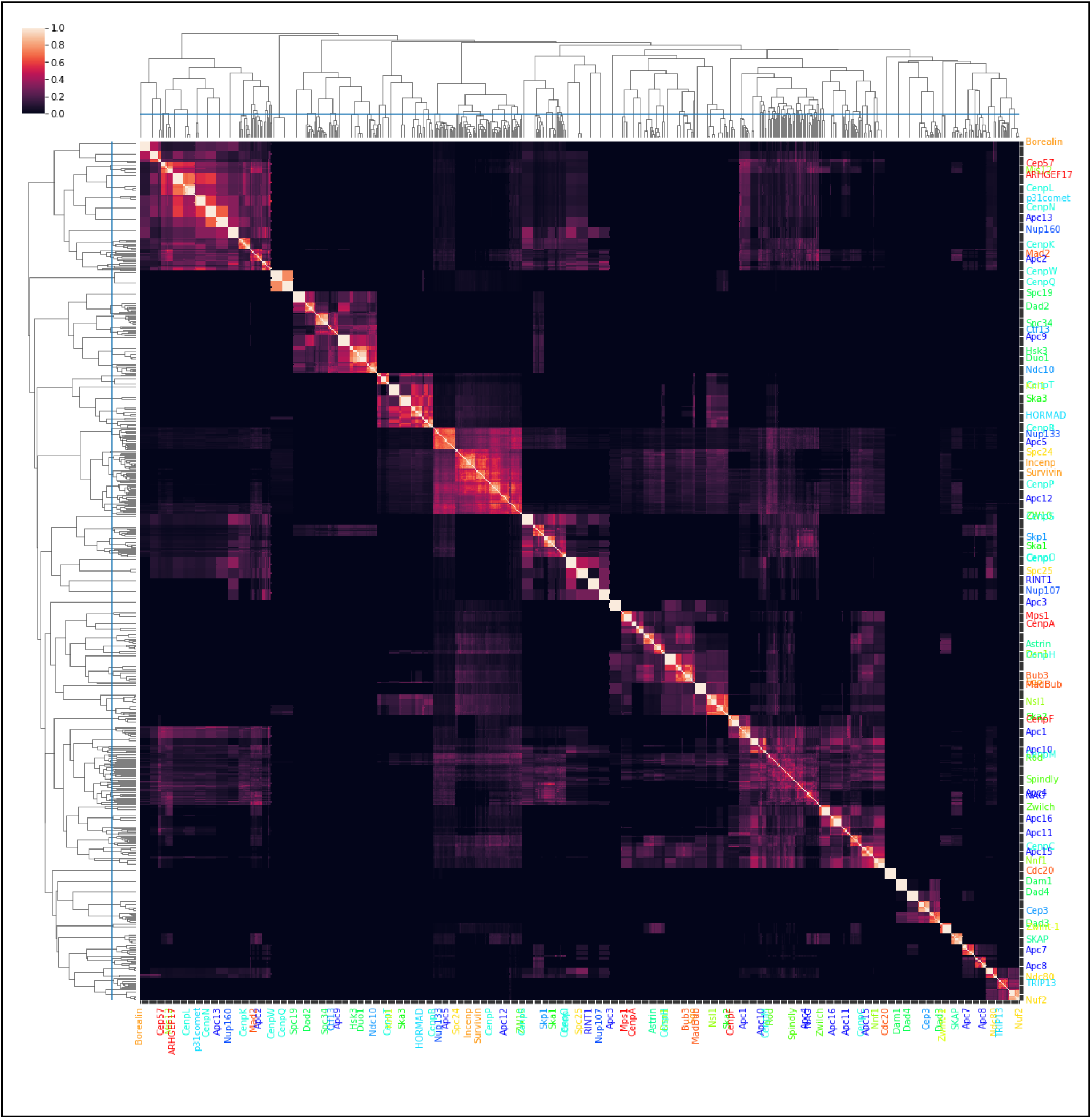
Putative novel components of the kinetochore and APC complexes. The profiles associated with all HOGs mapping to known kinetochore components shown in Figure 3 were used to search the LSH Forest and retrieve the top 10 closest coevolving HOGs resulting in a list of 871 HOGs including the queries from the original complexes. The Jaccard distance matrix is shown between the hash signatures of all query and result HOGs. UPGMA clustering was applied to the distance matrix rows and columns. Labelled rows and columns correspond to profiles from the starting kinetochore dataset [9]. A cutoff hierarchical clustering distance of 1.3 (blue lines) was used to generate a total of 136 clusters of HOGs used for GO enrichment to identify functional modules. The coloring of the protein family names to the right and below the matrix is identical to the complex related coloring shown in Figure 3. All code used to construct this figure is available in the HogProf repository.

**Fig. 5.**
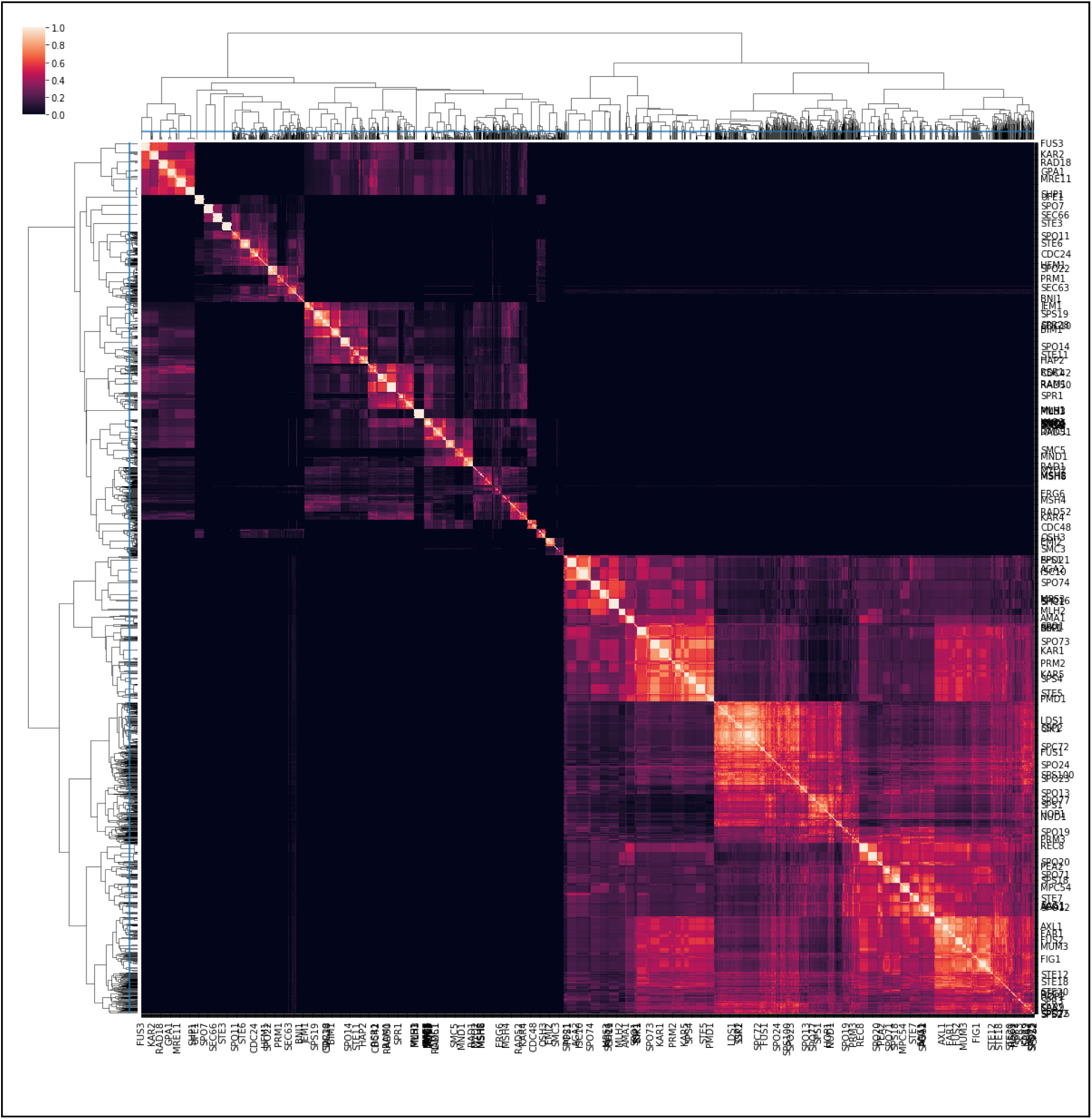
HogProf’s reproductive network. A list of proteins known to be involved in conserved sexual reproduction biological processes was compiled and each protein family was mapped to its HOG and used to search an LSH forest containing all HOGs in OMA. Each row and column of the Jaccard distance matrix corresponds to a HOG containing known sexual reproduction pathway protein families or a HOG returned by the search. A Jaccard distance matrix was generated by performing an All vs All comparison of the Hash signatures of the results and queries. UPGMA clustering was performed on the rows and columns to organize the HOGs into functional modules. The initial set of 121 protein sequences was augmented using the search functionality of the LSH by adding the top 20 closest returned HOGs resulting in a total of 2041 HOGs including the queries. A cutoff distance of .995 was used to generate a total of clusters of HOGs (blue lines). The labels correspond to the names of the proteins used to generate the queries. HOG names shown correspond to the yeast gene names (apart from Hap2 which is not present in fungi). This nomenclature was chosen due to the large body of work related to the yeast pheromone response and mating pathways. The HOGs returned by our search are not labelled on the distance matrix. All code used to construct this figures is available on the HogProf repository.

For instance, TACC3, a known physical interactor of the kinetochore complex and important regulator of the kinetochore tension [30]was found by our search. Another example is CFAP157, a cilia and flagella associated protein which may seem like an unlikely interactor with the APC. However, it has previously been shown that the APC activity regulates ciliary length unstabilizing axonemal microtubules [31]. Thus, CFAP157 might be involved in recruitment of APC regulators (such as Cdc20) or ciliary kinases (such as Nek1) both known to mediate APC regulation of ciliary dynamics [31]. While these results are certainly promising, many of the unannotated proteins returned by our search likely contain more regulatory, metabolic and physical interactors which may prove to be interesting experimental targets. The diverse types of interactions detected by our pipeline are discussed further in the discussion section.

### Search for a novel network

When studying networks with a lack of annotation and experimental characterization, it is difficult to quantify the relevance of retrieved search results. In typical research cases involving uncharacterized protein families in poorly studied neworks, this will often be the case. In this section we present the search results for three HOG queries known to be involved in the processes of meiosis, syngamy and karyogamy. These three major events occur in almost all sexually reproducing eukaryotes during their reproductive cycle. Despite the ubiquitous nature of sex and its probable presence in LECA [32], the protein networks involved in each part of these processes are very poorly understood and limited experimental data is available even in model organisms. However, some key protein families involved in these biological processes are known to have evolutionary patterns indicating an ancestral sequence in the LECA with subsequent modifications and losses [32]. The three following sections detail the returned results of the phylogenetic profiling pipeline with the Hap2, Gex1 and Spo11 families which all share this evolutionary pattern and are known to be critical for the process of gamete fusion, nuclear fusion and meiotic recombination, respectively. As in section 3.2 we also used GO enrichment to quantify the relevance of the returned search results. The proteins contained in the top 100 HOGs returned by the LSH Forest were analyzed for GO enrichment using all OMA annotations as a background. Due to the presence of biases in the GO annotation corpus [33] we have also chosen to show the number of proteins annotated with each biological process selected from the enrichment out of the total number of annotated proteins.

#### Query with Hap2

The Hap2 protein family has been shown to catalyze gamete membrane fusion in many eukaryotic clades [34,35]. It has a particularly spotty pattern of presence and absence on the taxonomic tree despite its phylogeny supporting the hypothesis of vertical descent from LECA. This protein family is known to be highly divergent in amino acid sequence despite its conserved fold and shares homology with viral and somatic membrane fusion proteins [35–37]. The HOG containing Hap2 in OMA only contains the eukaryotic gamete fusion protein subfamily of this structural superfamily. Part of the GO enrichment of the search results for the top 100 coevolving HOGs are shown below in table 3.

**Table 3.** Manually curated biologically relevant enriched GO terms from returned results. The chosen input protein sequence for Hap2 is that of UniProt entry F4JP36 and the corresponding OMA entry ARATH26614 belonging to OMA HOG:0406399. The full enrichment results are available in the Supplementary Data 2.

| Term | Biological process | P-value | N-proteins |
| --- | --- | --- | --- |
| GO:0006338 | chromatin remodeling | 9.72e-54 | 61/3426 |
| GO:0048653 | anther development | 1.69e-35 | 17/3426 |
| GO:0009793 | embryo development ending in seed dormancy | 2.88e-13 | 15/3426 |
| GO:0051301 | cell division | 6.88e-16 | 5/3426 |

Widely conserved sequences not belonging to the Hap2 HOG and found in coevolving HOGs were linked to gamete development and reproductive structure development (Table 3) [38,39]. This mirrors the initial discovery of Hap2, which was first found in angiosperms and linked to pollen tube guidance before the double fertilization event. Since Mendel, extensive work has been carried out describing reproductive processes in plants. Therefore, it is expected that the corpus of available annotations would be biased for annotations related to plant reproductive processes. The Hap2 HOG also appears to be coevolving with HOGs related to chromatin remodeling, an important part of the reproductive process during gamete generation, and also post fusion, after the zygote cell is formed.

One particular family of interest which was returned in our search results is already characterized in angiosperms: LFR or leaf and flower related [40]. This protein family is required for the development of reproductive structures in flowers and serves as a master regulator of the expression of many reproduction related genes, but its role in lower eukaryotes remains undescribed despite its broad evolutionary conservation. Experiments targeting LFR’s potential regulation of Hap2 expression may provide insight into how the fusion process is transcriptionally controlled in gametes across many eukaryotes despite their distinct reproductive strategies.

#### Query with Gex1

Gex1 has been shown to be involved in nuclear fusion (“karyogamy”) and is present in many of the same clades as Hap2, with a similar spotty pattern of absence across eukaryotes and a phylogeny indicating a vertical descent from LECA [41]. GO enrichment of the search results for the top 100 coevolving HOGs shows the predictive potential of HogProf (table 4).

**Table 4.** Manually curated biologically relevant enriched GO terms from returned results. The input protein sequence chosen for Gex1 is based on the UniProt identifier Q681K7 and the corresponding OMA identifier ARATH38809 belonging to OMA HOG:0416115. The full enrichment results are available as Supplementary Data 3.

| GO Term | P-value | N-Proteins |
| --- | --- | --- |
| GO:0042753 positive regulation of circadian rhythm | 2.12e-285 | 113/2685 |
| GO:0048364 root development | 7.81e-125 | 70/2685 |
| GO:0051726 regulation of cell cycle | 1.22e-92 | 99/2685 |
| GO:0000712 resolution of meiotic recombination intermediates | 1.65e-47 | 26/2685 |
| GO:0007140 male meiotic nuclear division | 1.19e-39 | 26/2685 |
| GO:0009553 embryo sac development | 1.43e-28 | 17/2685 |
| GO:0022619 generative cell differentiation | 3.59e-18 | 5/2685 |

In many sexually reproducing organisms, karyogamy is followed by restarting of the cell cycle. Our results strongly suggest that HOGs related to the restarting of the cell cycle have coevolved with Gex1 (Table 4). Again we find many angiosperm specific annotations due to the prior work in the study of their sexual reproduction. As was the case for Hap2, the taxonomic spread of the HOGs found in this search is broader than just angiosperms. Gex1 has also been shown to be involved in gamete development and embryogenesis [42] and therefore GO terms 0022619 and 0009553 are applied to this protein. Thus proteins that HogProf identified as putative Gex1 interactors sharing these GO terms indicates the potential relevance of these search results.

One result of particular interest is a protein family which goes by the lyrical name of parting dancers (PTD). PTD belongs to a family that has been characterized in Arabidopsis thaliana and yeast, and is known to be required in reciprocal homologous recombination in meiosis and localizes to the nucleus [43]. Our search shows that Gex1 coevolved closely with PTD, a protein known to be involved in preparing genetic material for its eventual merger with another cell’s nucleus.

#### Query with Spo11

Spo11 is a helicase that has been shown to be involved in meiosis by catalyzing DNA double stranded breaks (DSBs) triggering homologous recombination. Spo11 is highly conserved throughout eukaryotes and homologues are present in almost all clades [44]. The GO enrichment of the search results for the top 100 coevolving HOGs are shown below in table 5.

**Table 5.** Manually curated biologically relevant enriched GO terms from returned results. The chosen input protein sequence for Spo11-1 is based on the UniProt identifier Q9M4A2 and the corresponding OMA identifier ARATH19148 belonging to OMA HOG:0605395. The full enrichment results are available in Supplementary Data 4.

| GO Term | P-value | N-Proteins |
| --- | --- | --- |
| GO:0000737 DNA catabolic process, endonucleolytic | 0.00e+00 | 415/20562 |
| GO:0043137 DNA replication, removal of RNA primer | 0.00e+00 | 353/20562 |
| GO:0006275 regulation of DNA replication | 0.00e+00 | 552/20562 |
| GO:0006302 double-strand break repair | 8.11e-242 | 285/20562 |
| GO:0007292 female gamete generation | 2.71e-184 | 136/20562 |
| GO:0022414 reproductive process | 1.66e-93 | 127/20562 |

The process of chromosome recombination is one of the crucial steps in the generation of gametes and happens during meiotic prophase I when homologous chromosomes are paired and form the synaptonemal complex. It is encouraging to find that Spo11, the trigger of meiotic DSBs, has coevolved with other families involved in the inverse process of repairing the DSBs and finishing the process of recombination (Table 5). Other identified HOGs contain annotations such as gamete generation and reproduction also focusing at processes that result in cellular commitment to a gamete cell fate through meiosis. Proliferating cell nuclear antigen or PCNA [45] was also retrieved by our search. This ubiquitous protein family is an auxiliary scaffold protein to the DNA polymerase and recruits other interactors to the polymerase complex to repair damaged DNA, making it an interesting candidate for a potential physical interactor with Spo11.

In summary, this HogProf search focused on three proteins involved in sexual reproduction yielded a list of promising candidate proteins.

#### A broader search for the reproductive network

A more in-depth treatment of the evolutionary conservation of gamete cell fate commitment and mating is available in previous publications [32,41,46–50]. Using these sources, a list of broadly conserved protein families known to be involved in sexual reproduction were compiled to be used as HOG queries to the LSH Forest to retrieve the top 10 closest coevolving HOGs. The hash signatures of the queries and results were compiled and used in an all-vs-all comparison to generate a Jaccard distance matrix.

The all-vs-all comparison of the Jaccard distances between these returned HOGs reveals clusters of putative interactors coevolving closely with specific parts of the sexual reproduction network. The GO enrichment of sequences within each cluster was analyzed manually and several annotations related to sexual reproduction were found. These are summarized in Table 6 after a manual curation and literature review as done in table 2 for the kinetochore search results. In addition to annotated protein sequences and HOGs, many unannotated, coevolving HOGs where found. Again, these may prove to be useful experimental targets to answer open questions on the mechanisms behind sexual reproduction.

**Table 6.** Manually curated biologically relevant putative interactors from sexual reproduction search results. Notable protein families (Result) within clusters containing query HOGs (Cluster) are listed with their pertinent annotation and literature. GO enrichment results of clusters that contained one or more queries from our list of queries were analyzed manually. We searched for literature associated to the relevant GO annotations confirming results returned by the LSH that were associated with sexual reproduction. This is a non-exhaustive summary of some salient returned results. The full enrichment results are available in the Supplementary Data 5.

| Cluster | Result | GO Term | Citation |
| --- | --- | --- | --- |
| REC8 | NSE4 | GO:0030915 Smc5-Smc6 complex | [51] |
| SPC72 | MID2 | GO:0000767 cell morphogenesis<br>involved in conjugation | [52] |
| SPO71 | LES2 | GO:0031011 Ino80 complex | [53,54] |
| SHC1, SPO16 | POG1 | GO:0000321 re-entry into mitotic cell<br>cycle after pheromone arrest | [55,56] |

Particularly for biological processes as complex and evolutionarily diverse as sexual reproduction, Gene Ontology annotations are, unsurprisingly, incomplete. Fortunately, our profiling approach is successful in identifying protein families with similar evolutionary patterns that have already been characterised and are directly relevant to sexual reproduction (Table 6). By considering the uncharacterized or poorly characterized families at the sequence and structure level, we may be able to predict their functions and reconstitute their local interactome. Our ultimate goal is to guide *in vivo* experiments to test and characterize these targets within the broader context of eukaryotic sexual reproduction.

This example related to the ancestral sexual reproduction network illustrates the utility of the LSH Forest search functionality and OMA resources in exploratory characterization of poorly described networks. The interactions presented above in Table 6 only represent our limited effort to manually review literature to highlight potentially credible interactions detected by our pipeline. Again, as was the case with our kinetochore and APC related searches, several interactions might not appear obvious on their face. For example, SPC72 and MID2 are both involved in meiotic processes but localized to different parts of the cell (the centriole and the membrane respectively). However, it has been shown that microtubule organization and membrane integrity sensing pathways do show interaction during gamete maturation [57]. Others, like the Ino80 complex related Les2 subunit and SPO71 appear to be directly involved in the biological process of DNA remodelling during recombination and it may be easier to imagine their mode of interaction and design experiments to probe it.

## Discussion

We introduced a scalable system for phylogenetic profiling from hierarchical orthologous groups. The ROC and AUC values shown in the benchmark of section 3.1 indicates that the minhash Jaccard score estimate between profiles is a competitive alternative to previous tree and vector based metrics, while also being much faster to compute. This is remarkable in that one typically expect a trade-off between speed and accuracy, which does not appear to be the case here. We hypothesise that the error introduced by the fast minhash approximation is more than compensated by the inclusion of an unprecedented amount of genomes and taxonomic nodes in the labelled phylogenies used to construct the profiles.

Furthermore, while our minhash-derived Jaccard estimates are able to capture some of the differences between interacting and non-interacting HOGs, as shown above, their unique strength lies in the fast recovery of close profiles. Once these profiles are recovered, the inference of submodules or network structure can be refined using other, potentially more compute intensive methods, on this much smaller subset of data.

Because phylogenetic profiling is not yet broadly used on eukaryotic data, HogProf is largely orthogonal to and thus particularly effective combined with existing functional annotations. We showed that HogProf was able to reconstitute the modular organisation of the kinetochore, as well as increase the list of protein families interacting within the network with several known interactors of the kinetochore and the APC. As for the other HOGs returned in these searches, our results suggest that some are yet unknown interactors involved in aspects of the cell cycle or ciliary dynamics. Likewise, our attempt at retrieving candidate members of the sexual reproduction network recapitulated many known interactions, while also suggesting new ones.

The current paradigm for exploring interaction or participation in different biological pathways across protein families relies heavily on data integration strategies that take into account heterogenous high-throughput experiments and knowledge found in the literature. Many times, these datasets only describe the networks in question in one organism at a time. Furthermore, signaling, metabolic and physical interaction networks are all covered by different types of experiments and data produced by these systems is located in heterogeneous databases. By contrast, phylogenetic profiles can potentially uncover all three types of networks from sequencing data alone. This was highlighted in our work during retrieval of potential interactors within the sexual reproduction and kinetochore networks with the retrieval of LFR and CFAP157, respectively. In both cases, a regulatory action within the network was the biological process which involved both the query and retrieved HOGs, not a physical interaction. The advances put forward by our new methodology and the property of retrieving entire networks and not just physical interactions opens the possibility of performing comparative profiling on an unprecedented scale and lays the groundwork for integrative modeling of the interplay between PPI, regulation and metabolic networks in a more holistic way.

Further work remains to be done on tuning the profile construction with the appropriate weights at each taxonomic level, as well as when to construct profiles for subfamiles arising from duplications which may undergo neofunctionalization. Downstream processing of the explicit representation of the data, as opposed to the the hash signature, can also be designed using more computationally intensive methods to detect interactions on smaller subsets of profiles after using the LSH as a first search.

The phylogenetic profiling pipeline presented in this work will be integrated into OMA web-based services. Meanwhile, it is already available on Github as a standalone package.

## Methods

The following section details the creation of phylogenetic profiles using OMA data, their transformation into minhash based probabilistic data structures and the technical details of the implementation.

### Profile construction

To generate large-scale gene phylogenies labelled with speciation, duplication and loss events (a.k.a. *enhanced phylogenies* or *tree profiles*) for each HOG in OMA, we processed input data in OrthoXML format [58] with pyHam [59], using the NCBI taxonomic tree [60] pruned to contain only the genomes represented in OMA [15]. Tree profiles contain a species tree annotated at each taxonomic level with information on when the last common ancestor gene appeared, where losses and duplications occurred and the copy number of the gene at each taxonomic level. More information on the pyHam inference of evolutionary events can be found in [59].

Using this gene tree representation of the HOG, a multiset for presence, loss and duplication at each taxonomic level is compiled into a vector representation. In this representation each column corresponds to an evolutionary event or presence of a gene at a specific taxonomic level and the weight in the column corresponds to the weight (or importance) given to each node of the taxonomic tree for that class of events (Fig. 1). In this formulation, the Jaccard score between multisets [61] representing profiles will be more heavily influenced by nodes with a higher weight. In this manuscript only profiles with binary vectors are considered; the optimization of weighting and other refinements of the profiling pipeline will be the subject of future publications.

### Profile construction with Weighted Minhashing and Database construction using LSH Forest

Historically, distance metrics between profiles have fallen into two categories: tree-based and vector-based metrics [6,17]. Comparing all-vs-all profiles to define a distance matrix using metrics detailed in other phylogenetic profiling approaches, such as mutual information, Hamming distance or tree-aware methods [6,18,62–64], scales quadratically with the number of profiles. The time it takes to calculate profiles and a distance between two profiles typically scales poorly with the number of genomes considered, especially with tree-based methods. These computations are not practical when comparing the labelled phylogenies produced by pyHam for all HOGs in OMA, even with high performance computing.

Several studies have established the Jaccard similarity [65] between two profiles of presence and absence patterns across extant genomes as a valid phylogenetic profiling distance metric, which is able to capture an evolutionary signal closely related to shared protein functions [18,66,67]. This profile distance metric integrates well with the available algorithms and data structures available in the Datasketch library [68]. These data structures are built around minhashing techniques to retrieve similar sets of elements in sublinear time and allow a user to efficiently search the profile space without explicitly calculating the distance matrix between all profiles, as well as approximate the Jaccard similarity between profiles, by comparing hash signatures. Using these data structures to represent and search for the phylogenetic profiles effectively removes the necessity for an all-vs-all comparison.

Minhashing techniques were devised to measure the similarity of documents and search for similar documents within large datasets containing billions of elements [69–71]. A document can either be encoded as a set of unique words that occur within it or as a multiset representing the number of occurrences of each unique word. When dealing with sets where the total number of unique elements is unknown before processing all sets, it is preferable to encode them using the minhash algorithm, which allows the hash signature of the set to be updated as new unique elements are added without prior knowledge of all possible elements. When the total set of unique elements (e.g. all of the possible words in a corpus of documents or all of the taxa present in the species tree) is known, it is possible to use a minhash signature to represent the number of times each type of element occurs in a multiset of all possible elements. This representation is known as a weighted minhash and, depending on the dataset, may be more precise in retrieving relevant hash signatures (e.g. a document that mentions a specific word many times). The mathematical principles underpinning weighted minhashing and locality sensitive hashing forest algorithms and their implementation are described in earlier papers [61,72,73].

After transforming HOG profile vectors to their corresponding weighted minhashes using the datasketch library, an estimation of the Jaccard distance between profiles can be obtained by calculating the Hamming distance between their hash signatures [61]. The speed of comparison and lower bound for accuracy of the estimation of the Jaccard score is set by the number of hashing functions. The comparison of hash signatures has *O*(*N*) time complexity where N is the number of hash functions used to generate the minhash signature. Due to this property, an arbitrary number of elements can be encoded in this signature without slowing down comparisons. In our use case, this enables the use of an arbitrarily large number of taxa for which we can consider evolutionary events. With other metrics, such as Pearson correlation between vectors, the profile comparison between vectors scales linearly with the number or genomes or taxa considered in the best case scenario. In more complicated tree-based methods, these comparisons can be much more costly.

Weighted minhash objects can also be used to compile a searchable data structure referred to as a Locality Sensitive Hashing Forest (LSH Forest) [72]. The LSH Forest can be queried with a hash signature to retrieve the K neighbors with the highest Jaccard similarity to the query hash. The K closest hashes are retrieved from a B-Tree data structure [74]. This branching tree data structure allows for the dynamic insertion, deletion and querying of the LSH Forest data structure built upon it at orders of magnitude faster than previous profiling efforts. As previously mentioned, calculating linkages between all groups in non-probabilistic data structures requires an all-vs-all comparison of profiles which scales quadratically with the number of profiles in the dataset and can easily become computationally prohibitive. This penalty also applies whenever new genomes or taxonomic levels are added to the input matrix and the linkages must be recalculated. In the case of the LSH Forest, hash signatures of HOGs containing the new genome can be deleted and replaced in the database with a time complexity that scales logarithmically with the number of HOGs in the dataset. In non-probabilistic data structures, whenever a new HOG is added to an existing input matrix and linkages are recalculated, the penalty is linearly proportional to the number of HOGs already in the dataset and the number of HOGs added whereas in the case of the LSH Forest, the time complexity scales logarithmically with the number of HOGs already in the dataset and linearly with the number of HOGs added. Query time complexity in typical profiling approaches is heavily penalized for the number of orthologous groups and genomes included in the analysis whereas HogProf is unaffected by the number of genomes included (since it is only dependent on the number of hash functions used to generate the weighted minhash signature of HOGs) and scales logarithmically with the number of HOGs added to the database.

### Orthology data and software libraries used

Our dataset contains approximately 600,000 HOGs computed from the 2,167 genomes in OMA (June 2018 release) The main computational bottleneck in our pipeline is the calculation of the labelled gene trees for each HOG using pyHam. Even with this computation, compiled LSH forest objects containing the hash signatures of all HOGs’ gene trees can be compiled in under 3 hours (with 10 CPUs but this can scale easily to more cores) with only 2.5 GB of RAM and queried extremely efficiently (an average of 0.01 seconds over 1000 queries against a database containing profiles for all HOGs in OMA on an Intel(R) Xeon(R) CPU E5530 @ 2.40 GHz and 2 GB of RAM to load the LSH database object into memory). This performance makes it possible to provide online search functionality, which we aim to release in an upcoming web-based version of the OMA browser. Meanwhile, the compiled profile database can be used for analysis on typical workstations (note that memory and CPU requirements will depend on the number of hash functions implemented in the construction of profiles and the filtering of the initial dataset to clades of interest to the user).

All gene ontology (GO) annotations (encompassing molecular functions, cellular locations, and biological processes) for HOGs contained in OMA were analyzed with GOATOOLS [75]. To calculate the enrichment of annotations, the results returned by the LSH Forest annotations for all protein sequences contained in the HOGs returned by the search were collected and the entire OMA annotation corpus was used as background.

HDF5 files were compiled with H5PY (ver. 2.9.0). Pandas (ver. 0.24.0) was used for data manipulation. Labelled phylogenies were manipulated with ete3 [76]. Datasketch (ver. 1.0.0) was used to compile weighted minhashes and LSH Forest data structures. Plots were generated using matplotlib (ver. 3.0.2). PyHam (ver 1.1.6) was used to calculate labelled phylogenies for the HOGs in OMA.

### Pearson and Spearman correlation comparison of distance matrices

Distance matrices between all pairs of profiles in the kinetochore and APC complex protein families defined in [9] were compared using the Spearman and Pearson statistical analysis functions from the the SciPy python package to verify the monotonicity of the scores between families.

## Acknowledgements

We thank Monique Zahn for helpful feedback on the manuscript. This work was funded by a grant by the Novartis Foundation for Medical-Biological Research (#17B111 to CD), by the Swiss National Science Foundation (Grant 183723 to CD), by the Swiss Leading House for the Latin American Region (to CD and PSA), and by the Agencia Nacional de Promoción Científica y Tecnológica (PICT-2017-0854 to PSA).

## Conflict of Interest

none declared.

